# Large herbivore impact on plant biomass along multiple resource gradients in the Serengeti

**DOI:** 10.1101/2021.09.15.460562

**Authors:** Neha Mohanbabu, Mark E. Ritchie

## Abstract

Herbivores form an important link in the transfer of energy within a food web and are strongly influenced by bottom-up trophic cascades. Current hypotheses suggest that herbivore consumption and impact on plants should scale positively with plant resource availability. However, depending on the effect of resources on plant quantity and quality, herbivore impact may vary with different types of resources. We test four alternative hypotheses for the relationship between plant biomass, herbivore impact on plant biomass, and plant resource gradients, each based on how resources might affect plant abundance and quality to herbivores. We measured plant biomass for four non-consecutive years in a long-term grazing exclosure experiment in the Serengeti National Park that includes seven sites that vary substantially in rainfall and soil and plant nitrogen (N) and phosphorus (P). Our data supported the hypothesis that herbivore impact is controlled by plant quality, in this case driven by plant P, as herbivore effects on biomass decreased with higher rainfall but increased with greater plant P, but not N content. To our knowledge, this is the first experimental study to indicate that wild mammalian herbivory is associated with P availability rather than N. Our results suggest that P, in addition to water and N, may play a more important role in driving trophic interactions in terrestrial systems than previously realized.

## Introduction

Growing uncertainties in global climate patterns (IPCC 2014) and increasing inputs of nitrogen and phosphorus due to human activities (Peñuelas et al. 2013) emphasize the importance of understanding how trophic interactions might be influenced by changes in resources governing primary production. Bottom-up trophic cascades may strongly influence herbivore abundance, consumption, and distribution. Much of the variation in terrestrial plant consumption by insect and mammalian herbivores, estimated to be 5-90% of annual aboveground production (McNaughton 1985, Crawley 1989, Cyr and Face 1993), has been attributed to plant resource availability and climate gradients (Hawkins and Porter 2003, Moreira et al. 2015, Zhang et al. 2016). These attributions largely assume that plant and herbivore growth are limited by the same resource(s), and predict that herbivore populations and impacts on plant biomass should increase together with resource availability (Ben-Shahar and Coe 1992, Griffin et al. 1998, Cebrian and Lartigue 2004). However, a growing body of research demonstrates that plant biomass and production are simultaneously limited by multiple resources, namely water, nitrogen (N), and phosphorus (P) (Harpole et al. 2007, Elser et al. 2007, Cleland and Harpole 2010, Harpole et al. 2011, Fay 2015, Eskelinen and Harrison 2015) while herbivores during plant growing seasons are generally considered to be limited by plant N. In addition, plant quality to herbivores, as plant tissue resource: carbon (Sterner and Elser 2002), is likely to increase with N and/or P supply in the soil but decrease with any resource that promotes only C assimilation (e.g. light, water, CO2) (Jarell and Beverly 1981, McLauchlan et al. 2010). There is a need to disentangle the separate influence of these different resources rather than consider them in aggregate as “nutrients” (Olff et al. 2002, Borer et al. 2014, Borer et al. 2020), but few studies have done so.

We propose four non-mutually exclusive, alternative hypotheses for variation in fenced and unfenced plant biomass, and the impact of herbivore consumption on plant biomass along separate gradients of rainfall, N and P availabilities. In fenced plots, in the absence of herbivory, steady-state plant biomass is likely to increase with elevated resource supply, although under certain conditions of colimitation the steady-state plant biomass may remain unchanged across resource gradients (Sperfeld et al. 2016). In contrast, increasing resource supply may drive faster plant growth in unfenced plots leading to compensation for herbivory by plants, essentially minimizing herbivore impact on biomass (hereafter called *plant compensation hypothesis*) (Fig. 1a) (McNaughton 1983, Strauss and Agrawal 1999, Frank et al. 2002, Ritchie 2014, Ritchie and Penner 2020). Alternatively, elevated resource supply may support higher herbivore densities (Fritz and Duncan 1994, Hunter et al. 1997, Siemann 1998, Gruner 2004), presumably via increased plant growth, which may reduce plant biomass in unfenced plots more strongly and result in a positive association between herbivore impact and resource supply (hereafter *herbivore density response hypothesis*) (Fig. 1b). This hypothesis predicts that herbivore impacts may increase in response to resources that otherwise do not limit herbivore growth, such as phosphorus (P) in terrestrial systems but the evidence is still sparse (but see Schade et al. 2003, Joern et al. 2012). Yet increases in P may stimulate plant growth and foster a larger herbivore biomass (Bishop et al. 2010, La Pierre and Smith 2016) and impact on plant biomass (Mohanbabu and Ritchie, unpublished theoretical model). Few studies have explored whether mammalian herbivores and their effect on plant biomass are associated with P (Staver et al. 2021).

**Figure 1:**
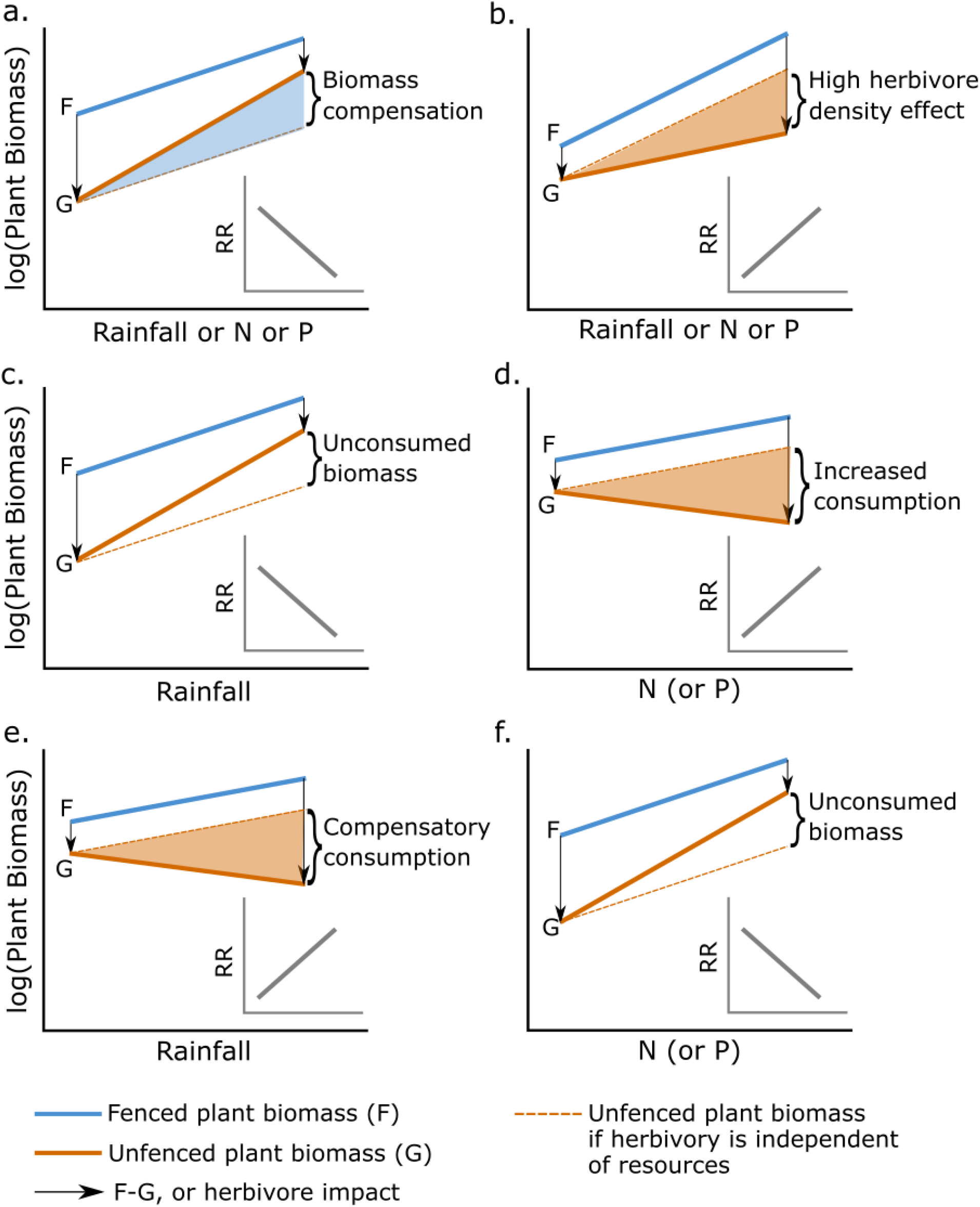
Non-mutually exclusive, alternate hypotheses for variation in fenced (F) and unfenced (G) plant biomass (log-transformed), and impact of herbivores on plant biomass (Response Ratio, RR) along gradients of rainfall, nitrogen, and phosphorus. a) Plant compensation hypothesis [increased plant regrowth reduces RR at high resource supply for ***<u>all</u>*** resources]; b) Herbivore density response hypothesis [increased herbivore densities in response to resources elevates RR at high supply of ***<u>all</u>*** resources]; c,d) Plant quality hypothesis [herbivores consume a lower proportion of plant biomass at high rainfall (i.e, low RR) but higher proportion at elevated N or P (i.e., high RR)]; and e,f) Herbivore compensation hypothesis [herbivores compensate for low quality by increasing consumption and RR at high rainfall and decrease consumption and RR at high N or P supplies]. The difference between the solid and dashed orange lines may be due to consumption replaced by plant regrowth (shaded blue), excess consumption at high resource supply (shaded orange), or unconsumed biomass (unshaded). Patterns across multiple resource gradients will be key in identifying the most likely hypothesis.

Increases in the supply of plant resources might also produce changes in herbivore impacts because they affect plant quality (resource: carbon stoichiometry, nutrient concentration, and the proportion of available biomass suitable for consumption) in different ways (hereafter *plant quality hypothesis*) (Fig 1c,d). The predominant hypothesis is that herbivores prefer plants with higher N as consumers have higher tissue nutrient content compared to that of producers (Sterner and Elser 2002). Under greater N availability, a greater proportion of plant biomass may be suitable for herbivore consumption (Mattson 1980, Schmitz 1994, Griffin et al. 1998, Welti et al. 2020), thereby causing a stronger reduction in plant biomass in unfenced plots. In contrast, the greater water supply may increase plant biomass without increasing plant N (Harpole et al. 2007, Cleland and Harpole 2010) and thereby may decrease plant tissue N (Luo et al. 2017, Wang et al. 2017, Welti et al. 2020) along with declines in the efficiency of herbivore growth per unit plant biomass, the proportion of palatable plant biomass and herbivore impact on plant biomass. Lastly, and alternatively, herbivores may also respond to poorer quality plants by consuming larger quantities of poor-quality plants to compensate for low nutrient content (Berner et al. 2005, Huberty and Denno 2006, Hillebrand et al. 2009, Couture et al. 2016). Under such a scenario, steady-state unfenced plant biomass would decrease with water supply and increase with N supply whereas herbivore impact would increase with greater water supply but decrease with higher N supply (hereafter *herbivore compensation hypothesis)* (Fig 1e,f). Associations of unfenced plant biomass and herbivore impact with P availability may not be strong given the lack of evidence for limitation of large mammals by P. However, if P supply determines the quality of plants, then responses of plant biomass and herbivore impact should resemble that of N supply (i.e, Fig 1d,f).

In this paper, we tested these alternate hypotheses simultaneously, and, to our knowledge, for the first time in terrestrial ecosystems. Even though exclosure studies are popular in grazing ecosystems, data from sites that span multiple resource gradients that also have a comparable herbivore species assemblage are very rare. The Long-Term Grazer Exclosure (LTGE) experiment in the Serengeti National Park, Tanzania (Anderson et al. 2007b, Ritchie 2014, Veldhuis et al. 2019a, 2019b) is unique as it spans natural gradients of rainfall, total soil N and P content, and plant N and P content while also being grazed by similar species of wild mammalian herbivores. Therefore, we measured within-year steady-state plant biomass and herbivore impacts on plant biomass, defined as the ratio of plant biomass in fenced to unfenced plots, at the LTGE experiment and analyzed data from four years to assess the effects of mammalian herbivores on plant biomass across multiple resource gradients.

## Study site and Methods

### Study area

The Serengeti National Park (SNP) in northern Tanzania is one of the last remaining intact grazing systems in the world, featuring more than 30 species of mammalian herbivores (Sinclair and Norton-Griffiths 1979, McNaughton 1985). However, these herbivores are heterogeneously distributed in hotspots (Anderson et al. 2010) and annually varying migration routes of wildebeest (*Connochaetes taurinus*) and plains zebra (*Equus quagga*), yielding substantial variation in forage consumption across the landscape. SNP also features large landscape gradients in rainfall and soil N and P (McNaughton 1985, Ruess and Seagle 1994, Anderson et al. 2007b), which may account for some of the variation in herbivore impact.

### Grazing exclosure experiment

Long-term grazing exclosures (LTGE) were established in 1999 at seven different sites separated by 20-100 km (Figure S1), along rainfall and nutrient gradients (Anderson et al. 2007b) (Table S1). Sites were chosen to be close to grazing hotspots of high densities of multiple resident herbivore species (Anderson et al. 2010), so LTGE sites are regularly grazed irrespective of annual wildebeest and zebra migration routes (Anderson et al. 2007b, Veldhuis et al. 2019b). As these hotspots are also associated with reduced risk of predation (Anderson et al. 2010), top-down control of herbivores is likely to be limited and therefore, unlikely to influence our results on herbivore impact on plant biomass. Mean annual rainfall ranges from 498-891mm, total soil N varies from 0.13 to 0.33%, and total soil P spans an order of magnitude from 0.007 to 0.08% (Anderson et al. 2007b)(Table S1). Each site features six 4 x 4 m plots arranged 30 m apart along a 180 m transect, three of which are fenced with chain link (mesh size ~10 x10 cm) at ~2m height and then reinforced with additional chain link (mesh size ~ 5 x 5 mm) at ~30 cm from the ground to exclude herbivores larger than 50 g and smaller than giraffes or elephants. Although the fences are not large enough to prevent giraffes and elephants, these mega-herbivores are likely to browse on woody plants which were virtually absent from the study sites (Anderson et al. 2007b) and never encountered in the clipped biomass. The relatively small size of the plots is also unlikely to support significant populations of small mammals, further corroborated by little evidence of small mammal activity inside exclosures (Anderson et al. 2007b). Nearest neighbor fenced and unfenced plots at each site were then paired to compare biomass for each of the three plot pairs at each site.

### Plant biomass

In each plot, we measured within-season steady-state plant biomass as the biomass at the end of the growing season (late May or June), as this best measures the season-long cumulative effects of consumption and plant regrowth. We clipped all plant material at the base in four randomly chosen 25 x 25 cm^2^ quadrants in each plot. Clipped material was sorted into live, brown litter (assumed to have been produced during the wet season) and gray litter (produced the previous wet season) and dried at 45°C for seven days. Biomass was sampled before any potential prescribed burns in four non-consecutive years (2001, 2006, 2009, and 2016), at all the sites except for Balanites site in 2016. Although the end of the season or peak biomass is a frequently used sampling protocol for exclosure studies (Borer et al. 2014, Veldhuis et al. 2019b, Wigley et al. 2020, Borer et al. 2020), there may be uncertainties associated with single time point measurements which was likely reduced by including data from multiple years.

### Site characteristics

We estimated the annual rainfall as the precipitation accumulated from the start of the dry season (July) of the previous year to the end of the wet season (June) of the year in which sampling occurred, based on the monthly averages from the CHIRPS database (Funk et al. 2015). Plant foliar N and P were measured from clipped aboveground green biomass (including flowers/ seeds if present), using the Kjeldahl method and persulfate digestion method, respectively, at the Sokoine University of Agriculture, in 2001 and 2016. Since we lack data on nutrient content for all the years, we used the measured values of plant N and P for 2001 and 2016 and values averaged over 2001 and 2016 years for 2006 and 2009. Soil N and P were measured for each plot during 2008 and 2016 using the same analysis methods. In statistical analyses, we used the values of soil nutrients averaged over both 2008 and 2016 for 2001 and 2006. Although plant N increased with soil N (Fig S2a), plant P showed considerable variation that was unexplained by soil P, especially at low soil P (Fig S2b). This mismatch suggests that plants may maintain tissue P despite low soil P as a result of mutualistic resource exchanges (Antoninka et al. 2015, Soka and Ritchie 2016, Ritchie and Raina 2016) or due to changing plant community composition (Ceulemans et al. 2014, Anderson et al. 2018). Regardless of the mechanism, herbivores ultimately respond to plant nutrient concentrations. Therefore, we also included plant N and P in our analyses as bioassays of the ability of plants to acquire nutrients, given N and P pools in organic matter and the dynamics of minerals mediated by plant-driven carbon supply and microbial activity.

### Statistical analyses

We define the impact of herbivores for each paired plot as a response ratio (RR) of plant biomass in the fenced to the unfenced plot (similar to Borer et al. 2020, Staver et al. 2021). All continuous variables, fenced and unfenced plant biomass, RR, and the predictors, were log-transformed for the analyses to meet the normality and heteroscedasticity assumptions of the linear models. We also checked all our models for multi-collinearity using Variance Inflation Factors (VIFs) and simplified the models to reduce multi-collinearity if VIF >10.

We used mixed models in R (R Core Team 2020) to test the effects of rainfall, N and P supplies on fenced and unfenced plant biomass, and herbivore impact on plant biomass (i.e., RR) on four years (2001, 2006, 2009 and 2016) of data. We fit linear mixed models with rainfall, total soil N and P, and plant N and P as the fixed effects and Site and Year as the crossed random effects, using the ‘blmer’ function from the blme package (Chung et al. 2013). The ‘blmer’ function uses a maximum penalized likelihood approach to estimate variance associated with random effects. Random effects with only a few levels, such as ‘Year’ in our case, are prone to ‘boundary estimate’ errors (or variance estimates of zero) in models that use a maximum likelihood approach. To overcome this issue, we use the blme package which specifies a weakly informative prior to the random effects covariance matrix based on a Wishart distribution, to provide non-zero estimates for random effects (Chung et al. 2013). We also analyzed the data by considering ‘Year’ as a fixed effect in addition to rainfall and nutrient parameters and the results were similar to the model with ‘Year’ as random effect (Table S2).

We considered several models that differed in their main and two-way interaction effects but always have ‘Site’ and ‘Year’ as crossed random effects. We selected the model with the lowest Akaike Information Criteria (AIC) as the most parsimonious model and used a Chi-squared test (likelihood ratio test) from the ‘anova’ function to confirm if the models were statistically different (*p* < 0.05) from each other. To assure ourselves that the trends are not an artefact arising from using soil and plant nutrient data from multiple years, we also analyzed data from 2016 for which we had the full complement of biomass and nutrient data and found that the overall patterns were consistent with the full data set (Table S3).

We then used the estimated slope values, i.e., positive or negative slopes, to test the directions of influence predicted by the hypotheses in Figure 1. For example, if all resources are positively associated with plant biomass or RR, then our data would support the “herbivore density hypothesis”. In contrast, if the unfenced plant biomass and RR are positively associated with some resources while being negatively associated with others, it may suggest that the “plant quality hypothesis” or the “herbivore compensation hypothesis” might better explain the data. For the purposes of visualizing associations of RR with different independent variables, we constructed partial residual plots (Fig 2) of variation in both dependent (RR) and independent (rainfall and N and P) variables once the covariance between independent variables is accounted for without any random effects. These plots matched the outcomes of the most likely linear mixed model with the random effects considered.

**Figure 2:**
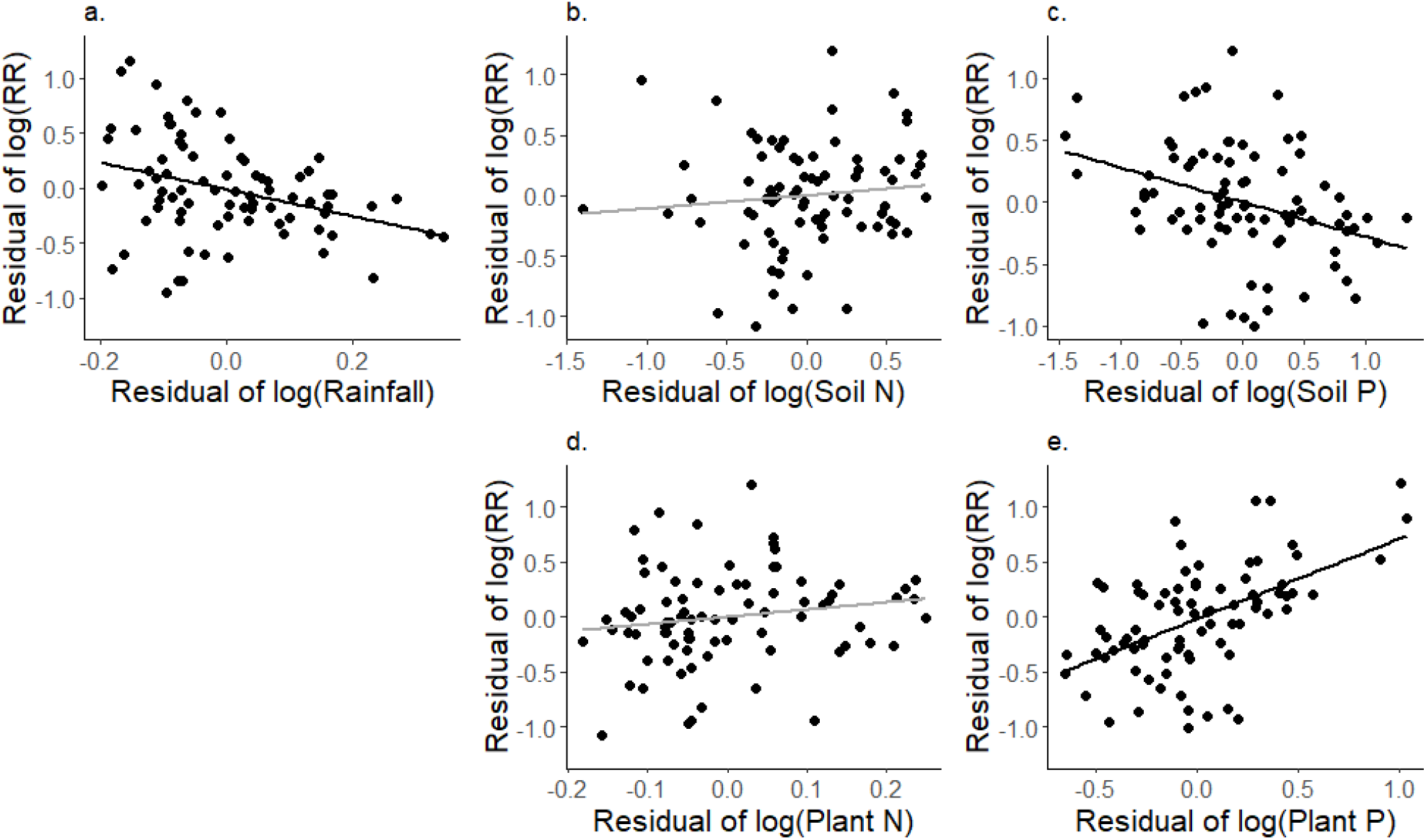
Variation in Response ratio (RR) along gradients of a) Rainfall; b) Soil N; c) Soil P; d) Plant N; and e) Plant P based on the most parsimonious statistical model i.e., the Phosphorus model. Both x and y axes are partial residuals to visualize the trend after accounting for other effects. Dark regression lines correspond to significant slopes (*P* < 0.05); light-colored lines indicate non-significant trends. See Methods section for details on the calculation of partial residuals.

## Results

The end of the season plant biomass in fenced plots were not associated with any of the resource gradients (Table 1a), as evident from the random intercept model being the most likely model. In contrast, the end of the season unfenced plot biomass increased with rainfall (Slope (SE): 1.15(0.61)), decreased with plant P (−0.73(0.28)), and remained unchanged with plant N, and total soil N and P when included in the model (Table 1b). In both cases, interaction terms resulted in high levels of multi-collinearity and had to be dropped from the models.

**Table 1:**
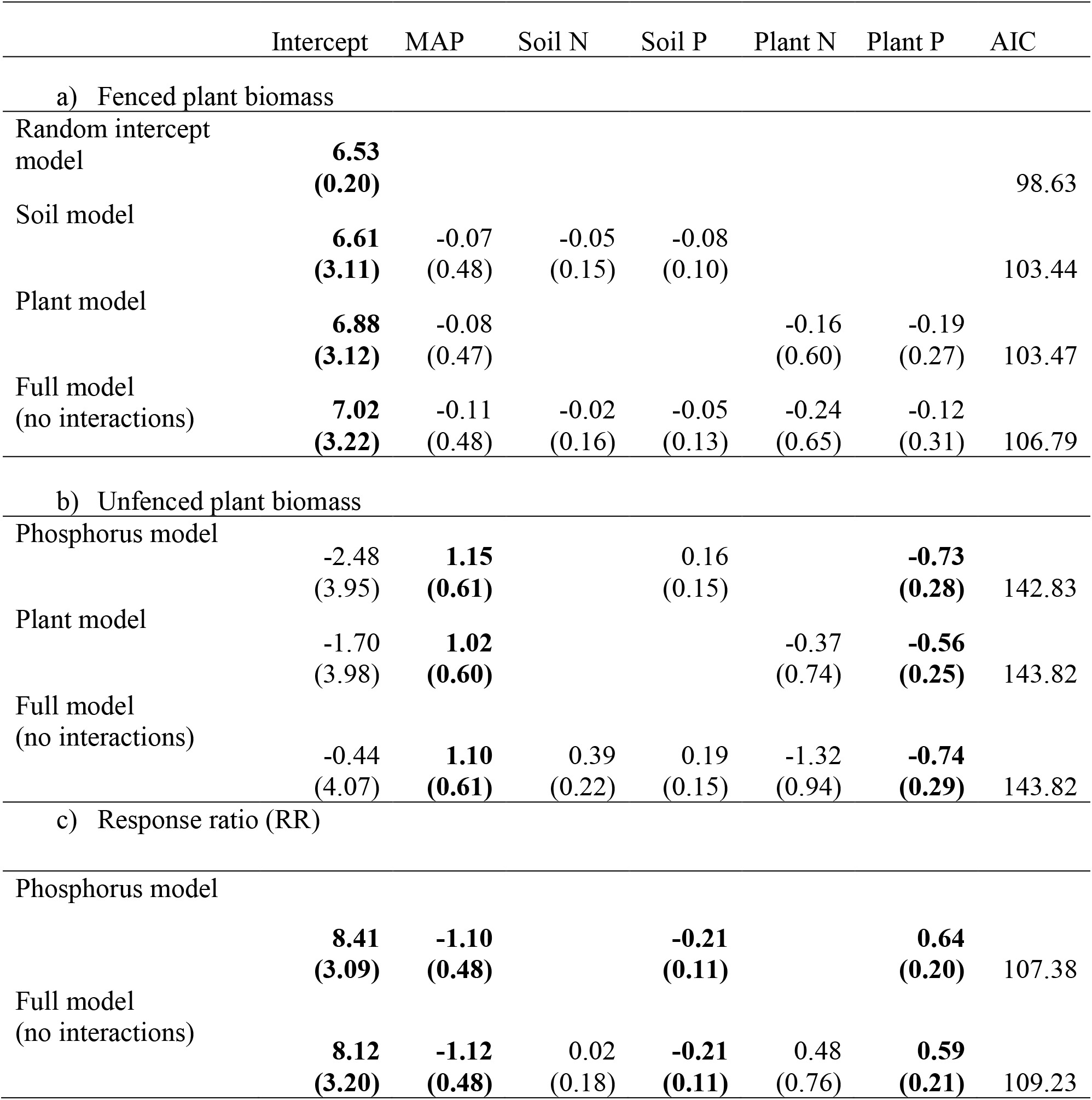
Estimates of AIC and slopes (SE in parentheses) associated with rainfall, soil and plant nutrients for the most likely models for a) Fenced plant biomass, b) Unfenced plant biomass and c) Response ratio or Herbivore impact. Estimates from other models that were not statistically different from the most likely model are also included. Values in bold are statistically significant.

The response ratio measuring the impact of fencing on biomass varied almost by an order of magnitude from near 1 (no impact) to over 10 across sites and years. The environmental variables explained a significant amount of variation in RR. Of the several models, the most likely model was one with rainfall, total soil P, and plant P as the fixed effects, without any interaction terms (Phosphorus model, Table 1c). The second most likely model which was not statistically different from the previous model, was the one with rainfall, soil N and P, and plant N and P as the fixed effects, without any interaction terms (Full model (no interactions), Table 1c). Yet again, interaction terms were insignificant and also resulted in high multi-collinearity and were dropped from the models. In both models, RR exhibited a significantly negative slope with rainfall (−1.10(0.48))(Table 1c, Fig 2a) and soil P (−0.21(0.11))(Table1c, Fig 2c), and a positive slope with plant P (0.64(0.20))(Table 1c, Fig 2e). Unexpectedly, the slope of RR was not significantly different from zero with either total soil or plant N content (Table 1c, Fig. 2b, e).

## Discussion

We show that the relative impacts of large mammal grazing on plant biomass, measured as a response ratio RR, can vary dramatically with RR decreasing with rainfall and soil P, increasing with plant P, and remaining unchanged with soil and plant N (Table 1c). These different associations with three resources that commonly limit plant biomass indicate that herbivore consumption, to the extent it affects biomass, may be driven by the influence of multiple resources on both plant growth and quality. The patterns of RR decreasing with rainfall and increasing with plant P support the “*plant quality hypothesis*”, under an assumption that herbivores are limited by plant P. To our knowledge, this is the first terrestrial study that has explicitly shown that effects of terrestrial mammalian herbivores on plant biomass are associated with P availability. Additionally, we find that the steady-state plant biomass in unfenced rather than fenced plots varied significantly across resource gradients, suggesting that plant biomass (in the absence of herbivory) may be co-limited by multiple resources, and it further interacts with herbivory to drive patterns in unfenced plant biomass and RR. These findings underscore the need to consider multiple resources for understanding how trophic interactions might be influenced by environmental variables, especially those like water, N, and P that are subject to ongoing climate change and anthropogenic manipulation of global N and P cycles (Peñuelas et al. 2013, IPCC 2014).

Our data are consistent with the *plant quality hypothesis*, where plant quality is presumably driven by plant P. This outcome is somewhat surprising since most previous studies suggest that mammalian herbivore growth is limited by N (Mattson 1980, Schmitz 1994, Griffin et al. 1998), sodium (McNaughton 1988, Kaspari 2020), or calcium (Ca) (Grunes and Welch 1989, Mládková et al. 2018) but not P (but see Murray 1995, Moen and Pastor 1998 for theoretical expectations). However, relatively few studies have considered variation in dietary P independently of other variables in mammalian growth and reproductive studies (Dykes et al. 2018). In addition to being a key component of DNA and RNA, P is also important for bone development in mammals (Grasman and Hellgren 1993). Thus, it is not unreasonable that P might limit herbivore growth and that herbivore impacts on plant biomass might increase with plant P. As expected from previous studies, herbivore impact on plant biomass declined with increasing rainfall, consistent with a mechanism by which plant biomass accumulation supported by high water availability “dilutes” nutrient content (Wang et al. 2017), leading to lower plant tissue P. The association with P may also be due to a herbivore density response to faster plant growth (quantities we did not measure) under greater soil P availability rather than a direct limitation of herbivore growth by plant P. However, the lack of association of fenced plant biomass with resources, but positive and negative association of unfenced plant biomass with rainfall and plant P, respectively, suggests that plant quality rather than quantity may drive these patterns in the Serengeti. It is possible that the historical emphasis on N as the key resource for plant-herbivore interactions (Mattson 1980, Bryant et al. 1983, McNaughton et al. 1997, Ritchie 2000, Holdo et al. 2007), in addition to relatively difficult methods for P estimation may have led to P being overlooked as an important resource for plant-herbivore interactions. Nonetheless, as evidence accumulates in favor of P (Schade et al. 2003, Bishop et al. 2010, Joern et al. 2012, La Pierre and Smith 2016) from different systems (temperate and tropical regions, mammal and insect herbivores), more work will be needed to elucidate the different mechanisms by which P might influence herbivore impact.

Support was generally lacking for the alternative hypotheses (Figure 1). The decline in RR with increasing rainfall does not support the *herbivore density response hypothesis* that higher rainfall yields faster plant growth, higher herbivore abundance, and greater herbivore impacts. The decline in RR with increasing rainfall also does not support the *herbivore compensation hypothesis* that impacts on biomass would be higher if plant quality was lower, presumably at higher rainfall. Furthermore, our data do not support the *plant compensation hypothesis* that herbivore impacts would be reduced at higher resource availability, which would support greater plant compensation to herbivory events at high resource availabilities (McNaughton 1983, Frank et al. 2002). Although our finding shows a negative association between RR and rainfall and soil P, which would support the plant compensation hypothesis, we found positive, not negative associations between RR and plant P. Additionally, both unfenced and fenced plant biomasses are not associated with soil P, further negating the plant compensation hypothesis. However, the weak negative response of RR to soil P may arise from potential negative association between soil P and arbuscular mycorrhizal fungi that typically colonize grasses in the Serengeti (Soka and Ritchie 2016), which may enhance plant P (Smith and Read 2008) at low soil P, similar to other systems (Miller et al. 1995, Liu et al. 2012, Johnson et al. 2015, Propster and Johnson 2015).

Unexpectedly, we found no influence of N supply, as either soil or plant N, on plant biomass or herbivore impact, as would be expected under the plant quality hypothesis and presumed N limitation of herbivore growth (Mattson 1980). One possible explanation is, despite widely varying soil N of 0.05-0.3% (Ruess and Seagle 1994) across sites, the relatively narrow range of plant N (1.75% - 2.75%) encountered in our study may have weakened our ability to detect association of herbivore impacts with plant or soil N. Plant N was largely above a proposed threshold of 1.5% above which herbivores may select diets that balance other important nutrients (Behmer and Joern 2008, Felton et al. 2016). Some dominant grasses in the Serengeti such as *Themeda triandra*, *Digitaria macroblephara*, and *Pennisitum mezianum* exhibit root-associated nitrogen fixation (Ritchie and Raina 2016, Ritchie et al., in prep), which may allow plants to have greater N than can be expected from soil N alone.

Strikingly, relationships in RR and unfenced plant biomass along resource gradients were stronger with plant concentrations of P rather than soil P. Such a pattern may reflect the frequently observed disconnect between total soil nutrients and plant-available nutrients due to immobilization of nutrients in soil minerals and organic matter and dependence of N mineralization on plant carbon sources (Binkley and Vitousek 1989). Thus, plant tissue nutrients may be better indicators of nutrient supply to trophic interactions than soil measurements. Other resources, such as Na or silica (Si), that are not known to limit plant growth but can influence herbivore biomass intake and abundance (McNaughton 1988, Massey et al. 2009, Kaspari 2020), might have influenced our results if they were correlated with resources we did measure. However, previous surveys in Serengeti (Anderson et al. 2007a, McNaughton unpubl. data) suggest that soil and plant P are not strongly correlated with plant Na or Si and so such elements were unlikely to affect our results.

Compensation to herbivory by plants likely affected our biomass measurements in unfenced plots, as plots could have differed in the time since the most recent grazing events, with accompanying greater regrowth and lower RR (Mitchell and Wass 1996). However, time since the most recent grazing event would likely be longer at sites with lower herbivore density and fewer grazing events, implying that any reduction in RR would have been at sites with the lowest herbivore consumption. Therefore, plant compensation following herbivory is unlikely to affect the direction of relationships we detected but might account for some of the variation in RR not explained by our statistical model.

Furthermore, it is possible that the fenced plots acted as a refuge from predators and trampling for small mammals, thus increasing their activity in fenced plots as has been reported for other grazing exclosures in African grasslands (Keesing 1998, Keesing and Young 2014). If so, we may have underestimated plant biomass in the absence of grazing and hence also herbivore impact from large herbivores outside the fences. However, it is unlikely to change the broader patterns that we see in our study. If the small mammal population does not change with resource gradients, similar to the findings of Young et al. 2015, then the relationships of plant biomass and RR would remain the same. Alternatively, small mammals may be more likely to seek refuge at sites with high herbivory levels implying that we may have underestimated herbivore impact from large herbivores at high grazing intensity sites, which can only weaken the associations of RR with resources. Therefore, the associations we see are despite the effects of small mammal herbivory, as they would have likely been stronger in the absence of small mammals.

## Conclusion

We tested several competing hypotheses about the influence of multiple resources on herbivore impact on plant biomass and conclude that herbivore impact is influenced by multiple resources. Our results were consistent with the “plant quality hypothesis” that herbivore impacts would be higher in response to forage nutrient content, in this case P, and lower in response to higher rainfall. This pattern represents the first time, to our knowledge, that impact of large mammalian herbivory on plant biomass has been shown to be positively associated with plant P rather than plant or soil N. However, it is unclear whether this pattern implies that herbivore growth might be limited by P or whether the pattern results from indirect positive influences of P on plant growth and herbivore density. Nevertheless, our results highlight the need to consider multiple plant resources in the environment in determining the strength of trophic interactions and herbivory in particular. Given the projected variation in rainfall in response to climate change and increasing anthropogenic inputs of nitrogen and phosphorus into the ecosystems, it will be crucial to understand the interactive effects of resources on plant-herbivore interactions.

## Acknowledgments

We would like to thank Emilian Mayemba for field assistance and Jason Fridley for feedback on statistical analyses. Special thanks to TAWIRI, COSTECH, and TANAPA for allowing us to carry out research at Serengeti National Park. The study was supported by NSF grants DEB 0308486, 0543398, 0842230, and 1557085.

## Author contributions

MR collected all the data; NM analyzed the data and wrote the first draft of the manuscript. Both authors contributed significantly to revising the manuscript.

## Data availability statement

Raw data (Veldhuis et al. 2019a) are available from Dryad: 10.5061/DRYAD.B303788.

