## Supplementary information for "Large herbivore impact on plant biomass along multiple resource gradients in the Serengeti"

Table S1: Site characteristics of the Long Term Grazing Exclosure (LTGE) experiment. Average soil N and P values are means of data from the grazed plots and the values in the parentheses are standard deviation representing within site variation. Rainfall data is the mean annual precipitation for the sites adapted from Anderson et al. 2007.

| Site | Avg Soil N (%) | Avg Soil P (%) | Rainfall (mm) |
|---|---|---|---|
| Barafu (BRS) | 0.33 (0.030) | 0.08 (0.034) | 498 |
| Soit (SOT) | 0.15 (0.062) | 0.08 (0.046) | 538 |
| Musabi Plains (MSB) | 0.17 (0.075) | 0.05 (0.021) | 891 |
| Togoro Plains (TOG) | 0.13 (0.055) | 0.04 (0.026) | 676 |
| Balanites (BAL) | 0.19 (0.096) | 0.03 (0.004) | 711 |
| Klein's Camp West (KCW) | 0.19 (0.048) | 0.007 (0.001) | 766 |
| Kuka Hills (KUH) | 0.15 (0.051) | 0.007 (0.002) | 779 |

Table S2: Comparison of estimates of slope and associated standard errors (in parentheses) for the different resource gradients when “Year” is considered as a fixed or a random effect. The estimates are presented for the two most likely models in each case. Values in bold are statistically significant associations.

| Parameters | P model; Year random | P model; Year fixed | Model 2; Year random | Model 2; Year fixed |
|---|---|---|---|---|
| Rainfall | **-1.10 (0.48)** | **-1.19 (0.54)** | **-1.12(0.49)** | **-1.19(0.55)** |
| Total Soil P | **-0.21 (0.11)** | **-0.18(0.12)** | **-0.21(0.11)** | **-0.18(0.12)** |
| Plant P | **0.64 (0.20)** | **0.56 (0.24)** | **0.59(0.21)** | **0.51(0.24)** |
| Total Soil N | NA | NA | 0.02(0.18) | 0.03(0.19) |
| Plant N | NA | NA | 0.48(0.76) | 0.42(0.78) |

Note: The four models are in blmer syntax are:

P model; Year random:

log(RR)~log(map)+log(soilp)+log(plantp)+(1|Year)+(1|Site)

P model; Year fixed:

log(RR)~log(map)+log(soilp)+log(plantp)+Year+(1|Site)

Model2; Year random:

log(RR)~log(map)+log(soilp)+log(plantp)+log(soiln)+log(plantn)+(1|Year)+(1|Site)

Model2; Year fixed:

log(RR)~log(map)+log(soilp)+log(plantp)+log(soiln)+log(plantn)+Year+(1|Site)

Table S3: Summary of the mixed model for variation in log(RR) along multiple resource gradients for the year 2016. Values in bold are statistically significant ($p<0.05$) and values in italics are only marginally significant ($0.05<p<0.1$).

| Resource | Estimate (+/- SE) |
|---|---|
| log(MAP) | **-1.59(1.33)** |
| log(Total soil N) | -0.31(0.44) |
| log(Total soil P) | *-0.19(0.17)* |
| log( Plant N) | 0.59(0.99) |
| log( Plant P) | **0.55(0.40)** |

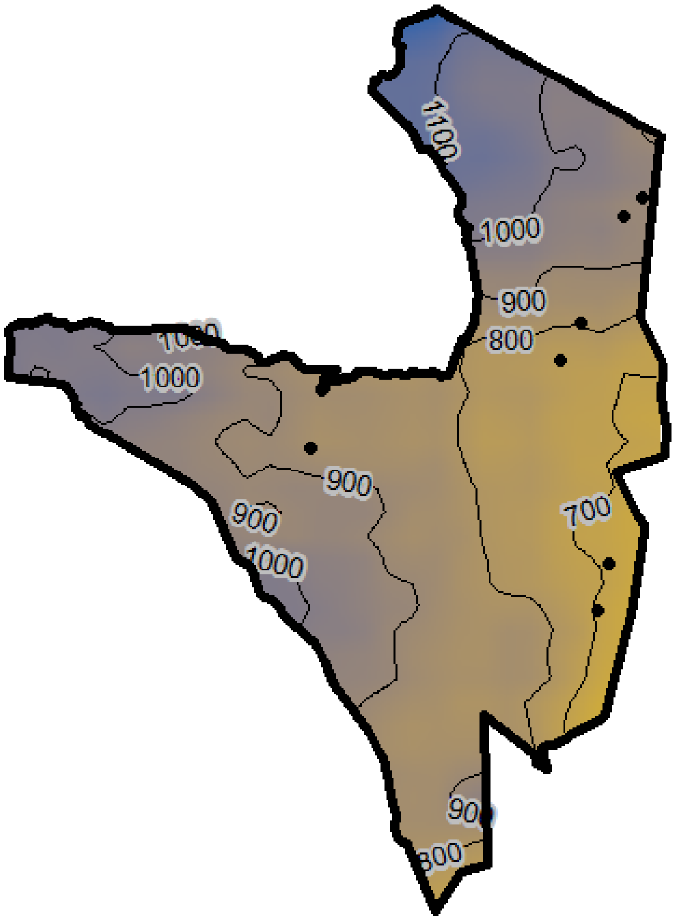


Fig S1: Sites that are being used for the Long Term Grazing Exclosure (LTGE) plots along with the contours for mean annual precipitation (Jan – Dec) from 2000-2016, based on CHIRPS data (different from the statistical model which uses data for the specific years from July- June). The site codes are BRS- Barafu, SOT- Soit, MSB- Musabi Plains, TOG- Togoro Plains, BAL- Balanites, KCW- Klien's Camp West, and KUH- Kuka Hills.

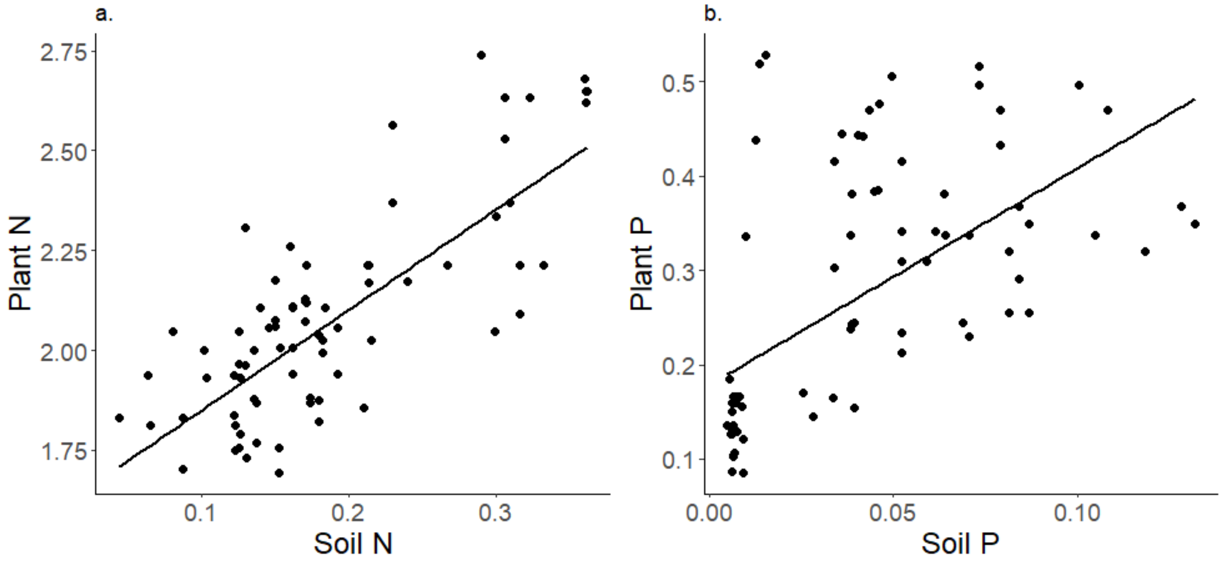


Figure S2: Association between soil and plant variables for a) Nitrogen; and b) Phosphorus. The slopes are both significantly different from 0, although soil N is a better predictor of plant N ($R^2$=0.57) than soil P is for plant P ($R^2$=0.32).
